# Spatial-temporal evolution characteristics of land use and its driving factor analysis during the period of 2000-2020 in Putian City, Fujian Province

**DOI:** 10.1101/2022.12.20.521249

**Authors:** Jinshan Zhang, Dongqing Wu, Qingxia Peng, Zhimin Lin, Wenxiong Lin, Kai Su

## Abstract

Exploring the spatial and temporal evolution characteristics of land use and its driving force can provide scientific basis for the construction of regional ecological civilization. Based on the remote sensing cloud computing platform of Google Earth Engine (GEE), the study took the remote sensing images of the three periods of 2000, 2010 and 2020 as the basic data and interpreted them to obtain the basic land use data of Putian city and its five districts and counties. The land-use change characteristics of Putian city in those periods were systematically analyzed by using the methods of single land-use dynamics and Geo-informatic Tupu, so as to reveal the Tupu characteristics of land-use transfer and the spatial-temporal evolution pattern in the past 20 years. Then social and economic development indicators were selected to further explore the key driving force of land-use evolution in the study area by canonical correspondence analysis. The results showed that: 1) the land use structure of Putian city was mainly composed of cultivated land and forest land, and the other land types were embedded in them and the built-up land continued to expand outward. 2) The spatial distribution of Tupu units of land use transfer in Putian city was significantly different, change types of land use were diversified and the areas of cultivated land and grassland transferred out were the most obvious. Different degrees of cultivated land abandonment had occurred in various regions. Putian city was facing great challenges in ensuring food security and curbing the non-agricultural and non grain of cultivated land. 3) The macro-economic development in a specific period, especially the urban expansion and the development of the secondary industry caused by the merger of cities and counties, were the key factors driving the spatial and temporal evolution of land use and the differential distribution pattern in Putian city. The author suggested that land resources should be used efficiently in addition to optimizing the layout of land use and carrying out the renovation of abandoned land. Therefore, the study proposed to strengthen the scientific planning and effective regulation of land use, combine spatial and temporal resources of cultivated land with agriculture, culture and tourism, and build ecological barriers and ecological corridors to ensure the coordinated and sustainable development of regional ecology and economy.

## Introduction

Land use / cover change (LUCC) is the most direct manifestation in the impact of human activities on the earth’s surface system, and has become one of important research areas in global environmental change and sustainable development. With the advancement of the rapid globalization, urbanization and industrialization in the world, especially in developing countries, the social and economic transformation has also had a significant impact on regional land use [1, 2], which makes land use transformation become a new theme and frontier of LUCC research.

Since the 1990s, China’s urban-rural transformation has led to a drastic land-use transition (LUT), which has greatly affected the ecosystem structure and its service functions, threatening the sustainable development of human beings and the sustainable supply of ecosystem services [3-5]. For example, the transformations of native grasslands, ecological forests and wetlands into farmland, artificial forests and built-up land, have greatly promoted a substantial increase in the production of food, wood, housing and other commodities, but paid a higher cost at a drastic reduction in many ecosystem services and biodiversity [6]. On the other hand, land use transformation is also an important driving factor of global climate change [7] and an important source of greenhouse gas (GHG) emissions. Some data show that agriculture, forestry, and other land use (AFOLU) activities cause about 20 GtCO_2_/year of anthropogenic greenhouse gas emissions, accounting for 20-24% of the total global emissions [8]. Therefore, in-depth study of the spatial and temporal characteristics of land use changes is not only conducive to clarify the process and causes of land use transformation, but also significant to the rational use of land resources and the sustainable development of social economy.

In recent years, the dynamic change of land use in time and space has been paid more and more attention by domestic and foreign researchers, and many research achievements have been obtained, which can provide important references for further exploring the process and causes of land use type transformation and improving resource utilization rate, and offer ideas for conducting relevant research in typical or specific areas. Researchers mainly choose the transformation of cultivated land, wetland and other land types [9, 10] as research objects and contents, and most of them pay more attention on the macro scale such as provinces, basins and plains [11-13]. Land-use dynamic index [14], land-use degree [15], land use comprehensive indexs [16] and other methods are used to analyze the spatial-temporal change characteristics of land use while spatial econometric regression analysis [17], landscape pattern index [18] and CLUE-S model [19] are adopted to study the evolution of land use spatial pattern. These above-mentioned research methods reveal the spatial and temporal evolution characteristics of land use transformation to a certain extent. However, the time process and spatial pattern of the land use transformation are not well combined, the characteristics thus obtained lack image sense, and the non-spatial data are not sufficiently expressed in terms of spatial location. The Geo-informational Tupu proposed by academician Chen Shupeng is a kind of spatiotemporal analysis method [20], which can present the spatial structure characteristics and temporal changes of objective things and phenomena through data mining and special processing, and generate a series of intuitive graphs, images and schema information in the form of Tupu units in the process of land use change [21, 22]. At present, some scholars have used such method to carry out research and initial achievements have been obtained. For example, Zhang *et al*. [23] explored the temporal and spatial characteristics of land use transformation in the Yellow River Basin by using Geo-information Tupu and rising/falling Tupu. Lu *et al*. [24] analyzed spatial transformation of the land in Shandong Province based on Geo-information Tupu and found that the land of Shandong Province was dominated by agricultural production space, and the expansion of urban living space was significantly higher than that of rural living space. In general, the existing literature mainly focuses on the statistical analysis and spatial-temporal distribution of land use Tupu, less on the reasons for the changes of Tupu units and the differences between regions and most of research areas are concentrated in river basins, economically developed areas and ecologically weak areas.

Coastal cities with abundant natural and economic resources are the first place where the marine economy started and their demand for sustainable development has brought new challenges to land use. Therefore, it is typical to explore the evolution of land use in coastal cities [25]. Putian is an emerging coastal tourist city in Fujian Province, rich in biodiversity resources. It is the national production base of longan, litchi and olive, the famous and excellent base of south subtropical crops (Loquat and Wendan pomelo) of the former Ministry of agriculture and has the reputation of “the first hometown of loquat in China”. As a key link in the construction of the Economic Zone on the West Bank of the Strait, Putian city plays an important role in the ecological security pattern of Fujian Province. In recent years, with the acceleration of highway/railway construction, industrialization and urbanization and other factors, a large number of agricultural land and ecological land in Putian city have been converted to built-up land, resulting in the contradiction between cultivated land protection, ecological conservation and urban expansion. Such resource consumption pattern has led to problems such as sensitive land use change and inefficient urbanization [26]. However, there are few studies on land use transformation and its ecological benefits in Putian city. Accordingly, it is necessary to monitor and analyze the change of land use, and timely explore the spatial and temporal evolution pattern of land use transformation and its causes in Putian city. Therefore, the study adopted the methods of Geo-informatics and canonical correspondence analysis to explore the spatial-temporal evolution pattern and its driving factor of land use in Putian city from 2000 to 2020, with a view to providing a decision-making reference for optimizing land resource allocation and its rational utilization, and hence formulating the sustainable development strategy of regional economy.

## Research area and methods

### Overview of the study area

Putian city is located in the central coastal area of Fujian Province (between 24°59’-25°46’ N and 118°27 ‘-119°39’ E), facing Taiwan Province across the sea. It has jurisdiction over five districts and counties, namely Licheng district, Hanjiang district, Chengxiang district, Xiuyu district and Xianyou county. It is the hometown of Mazu, the goddess of peace on the sea, and the world Mazu Cultural Center (Fig 1). The northwest of Putian city is a high-lying mountain range, and the southeast coast is mostly low hills and plains. It has Xinghua plain, the third largest plain in Fujian Province, which is the largest “land of fish and rice” in the city and belongs to marine subtropical monsoon climate, with an annual average temperature of 18°C - 21°C, abundant rainfall but uneven spatial distribution, and annual rainfall of 1000 - 1800 mm. The water system in Putian city is relatively developed, including Mulan River, Yanshou River and Qiulu River; the soil is mainly red soil and paddy soil; the vegetation is evergreen broad-leaved forest. In 2020, the forest coverage rate of the whole city reached 60.21%, and it won the title of “National Forest City”. Due to disturbance of production and construction projects, the soil erosion of slope farmland is relatively serious, and the same is true in the case under forests. In 2020, Putian’s GDP was 264.397 billion yuan, the resident population was 3.2107 million, and the proportion of primary, secondary and tertiary industries was 4.75%, 51.53% and 43.72% respectively.

**Fig 1.**
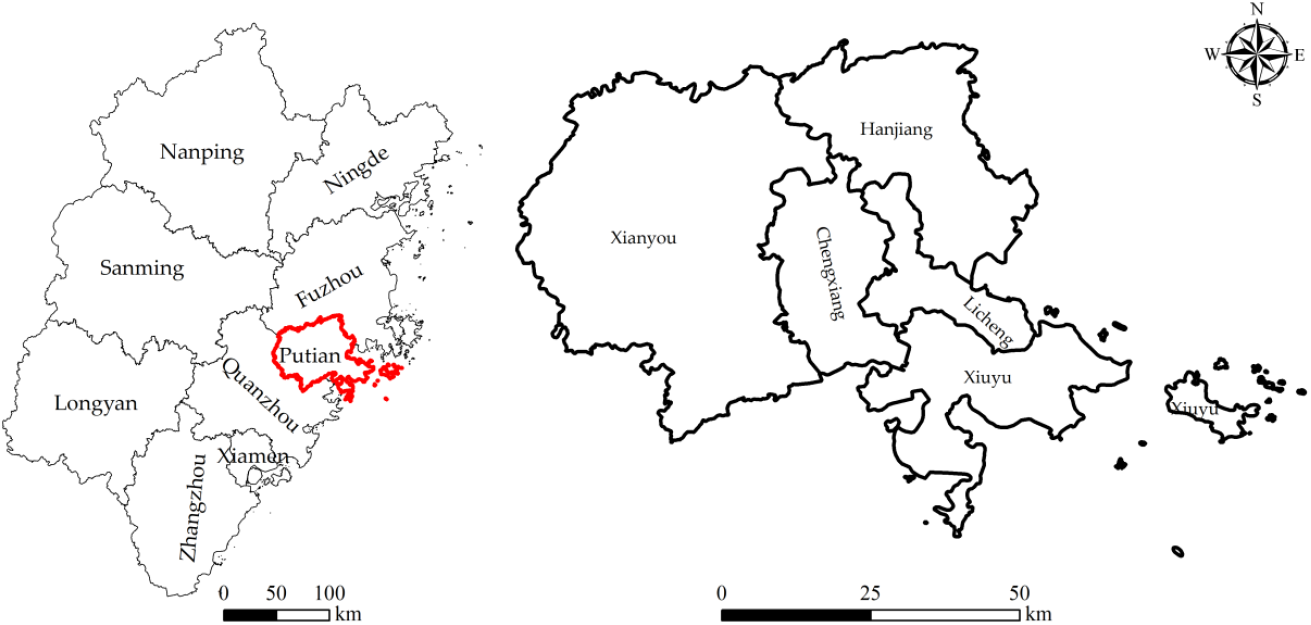
The location of study area.

### Data acquisition and processing

Based on the remote sensing cloud computing platform of Google Earth engine (GEE), the Landsat TM/OLI remote sensing images of the three periods of 2000, 2010 and 2020 with cloud cover less than 5% were selected as the basic data, with a spatial resolution of 30m. The normalized difference vegetation index (NDVI), normalized difference build-up index (NDBI) and enhanced vegetation index (EVI) were used as training variables of random forest (RF) algorithm to improve interpretation accuracy. According to *Current Land Use Classification* (GBT 21010-2017), the land use types were divided into cultivated land (CL), forest land (FL), orchard land (OL), grassland (GL), water body (WB), built-up land (BL) and unused land (UL) in combination with the actual conditions and research purposes of the study area. The land use spatial database of Putian city was established by setting the codes of each region as 1, 2, 3, 4, 5, 6 and 7 respectively. In this study, GEE, ArcGIS10.8 and other software were used as computing platforms.

## Research methods

### Random forest algorithm

Random forest algorithm (BF algorithm) is a supervised machine learning algorithm proposed by Leo Breiman in 2001 and widely used in classification and regression problems [27]. RF algorithm has high classification accuracy and has been widely used in land use classification by domestic and foreign scholars [28-30]. The basic principle of the RF algorithm is as follows: 1) use the Bootstrap sampling method, repeatedly and randomly extract *N* groups of training sets from the original data set and return them; 2) the training set of each group is about 2/3 of the original data, and the test set is about 1/3 of the original data; 3) each decision tree is composed of a group of corresponding training sets. During the construction of each tree, *m*(*m* ⩽ *M*) feature variables are randomly selected from *M* feature variables to divide the internal nodes. The subset composed of *m* variables is called feature random subspace; 4) integrate the prediction results of *N* decision trees and determine new sample categories by voting; 5) through the above steps, an RF model with *N* decision trees is established, its performance is scored by the test set, and the importance of each feature is ranked [31].

### Three spectral indices

NDVI is a dimensionless index describing the difference between visible light and near-infrared reflectance of vegetation coverage, which can be used to estimate the greening density on the land area. The calculation formula is as follows [32]:

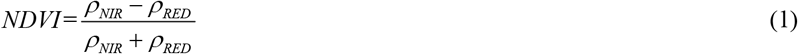

NDBI is used for automatic mapping of built-up areas. The main advantage of this index is its unique spectral response of built-up areas and other land cover. The calculation formula is as follows [33]

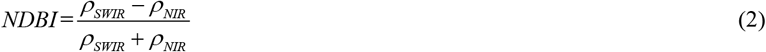

EVI is considered as an improved NDVI, which increases the sensitivity and vegetation monitoring ability in high biomass areas by decoupling the canopy background signal and reducing atmospheric impact. EVI was adopted by MODIS (modern resolution imaging spectroradiometer) terrestrial discipline group as the second global vegetation index to monitor terrestrial photosynthetic vegetation activities. The calculation formula is as follows [34]:

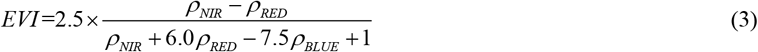

Where *ρ*_*NIR*_、*ρ*_*RED*_、*ρ*_*SWIR*_ and *ρ*_*BLUE*_ are the reflectance of near-infrared, red, short wave infrared and blue in Landsat images.

### Single land-use dynamic index

Single land-use dynamic index can quantitatively describe the speed of regional land use change, and play an important role in comparing the regional differences of land use transformation and analyzing the trend of land use change [4]. The formula is as follows:

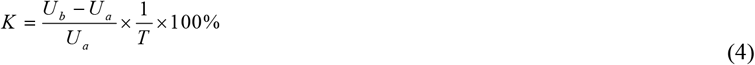

Where *K* refers to single land-use dynamic index, *U*_*a*_ and *U*_*b*_ represent the area of a certain land type at the beginning and the end of the study respectively, and *T* denotes the research period.

### Analysis of land use Tupu

Geo-informatic Tupu is a spatio-temporal analysis method that records land use change with Tupu units [20]. It has the dual properties of graph and pedigree. The graph and the pedigree are respectively used to represent spatial location characteristics and process variation [22]. The calculation formula of Tupu unit is as follows:

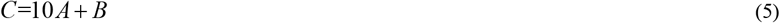

Where *C* represents the newly generated code value, that is, the type of land use Tupu unit; *A* and *B* refer to code values of the land use type at the previous stage and the later stage respectively. For example, code 21 means the conversion of cultivated land to forest land.

The transformation Tupu of land us in Putian city in 2000-2010 and 2010-2020 can be generated by comprehensively using the Tupu code fusion and map algebra superposition operation to synthesize the Tupu unit with the integration of characteristics of “space-attribute-process”. Land use transfer includes two aspects: transfer in and transfer out. The area transferred from other land types into this land type is classified as the new area of this land type, that is, the rising Tupu; all the areas transferred out of the land type are classified as the reduced area of the land type, that is, the falling Tupu. Therefore, the rising and the falling Tupu of Putian city in 2000-2010 and 2010-2020 can be obtained by using the spatial analysis of reclassification.

The transfer ratio of land use type [35] is used to calculate the proportion of a certain land type transfer to all land types transfer, which can further reflect the change characteristics of the number of LUT Tupu units. The calculation formula is as follows:

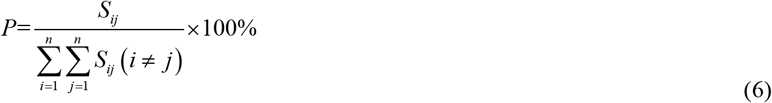

Where *P* represents the transfer ratio of land use type; *S*_*ij*_ refers to the area of Tupu unit of the *ith* land use type at the initial stage to the *jth* land use type at the final stage; *n* means the number of land use types.

### Accuracy evaluation model of land classification results

It is necessary to evaluate the accuracy of the interpretation results after applying classification algorithm to interpret remote sensing images. In this study, the overall accuracy (OA) and Kappa coefficient are used to evaluate the accuracy of land classification results [36], and the calculation formula is as follows:

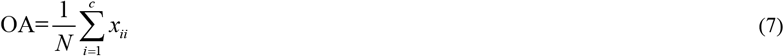

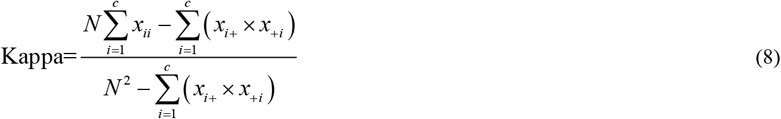

Where *N* and *x*_*ii*_ represent the total number of pixels and the total number of correctly classified pixels, respectively; *c* is the number of categories; *x*_*i+*_ and *x*_*+i*_ refer to the sum of rows and columns in the confusion matrix, respectively.

### Analysis on driving force of land use change

Canonical correspondence analysis (CCA) is a multivariate constrained ordination technique designed to extract comprehensive environmental gradients from eco-environmental datasets for elucidating the relationship between biological assemblages of species and their environment [37,38]. Therefore, this study uses the canonical correspondence method to quantitatively analyze key drivers of land use change.

The study selected seven types of land use in Putian city and its five districts and counties as species variables. According to the regional socio-economic development trend and data availability, 17 socio-economic indicators with extremely significant correlation with land use change were selected as the environmental variables, namely, fishery output value, the gross value of primary industry, GDP, total industrial output value, urbanization rate, the gross value of tertiary industry, the gross value of secondary industry, the area of economic crops, the non-agricultural population, the total retail sales of social consumer goods, the total population, and the agricultural population, which basically include the main socioeconomic indicators of the region.

CANOCO 4.5 and CANOD RAW 4.5 were used for CCA analysis.

## Results

### Analysis of land use structure change

The status map of land use classification in Putian City in 2000, 2010 and 2020 was obtained based on the GEE platform (Fig 1). According to formula (7) and formula (8), the overall accuracy of the three-phase remote sensing interpretation was 93.35%, 92.17% and 93.13% respectively, and their kappa coefficients were 88.13%, 86.64% and 87.51% respectively, suggesting that the interpretation results of the three-phase of land use classification had high accuracy and met the requirements of the research requirement.

As shown in Fig 2 and Table 1, forest land was the largest land type in Putian city, and the forest land area at the three-time courses exceeds more than 50% of the total area of the study area; the second was cultivated land, with an area of more than 1000 km^2^. The average ratio of the cultivated land area to the total area at the three-time courses reached 28.80%, indicating that the land use structure of Putian city in the past 2 decades was mainly based on agricultural and forestry production. However, the area of cultivated land, forest land and grassland decreased during this period, while the orchard land, water body, built-up land and unused land increased. Among them, cultivated land had the largest decline among all land types and showed a continuous downward trend, specifically from 1261.15 km^2^ in 2000 to 1003.29 km^2^ in 2020, with a decrease of 20.45%. Forest land was the land with the smallest area reduction among all land types, with a reduction of 24.79 km^2^ in the past 20 years. The grassland area decreased significantly from 2000 to 2010 with a reduction area of 45.44 km^2^ and an average annual reduction rate of 0.09% while it increased by 2.67 km^2^ from 2010 to 2020. As far as the orchard land was concerned, its area in 2000 was 12.76 km^2^, with an increase of 50.86% over the past 20 years, reaching 19.25 km^2^. The built-up land had increased from 109.15 km^2^ in 2000 (accounting for 2.78% of the total area) to 335.65 km^2^ in 2020, with an area of more than 2 times, making it the largest increase. The increase of unused land area was second only to that of built-up land, with an increase of 148.92% in the past 20 years. In addition, the water body area also showed a trend of continuous increase, which results from ecological restoration and governance project of Mulan River in this period.

**Table 1.** Statistics on the transformation of land use structure in Putian city from 2000 to 2020.

| Land type | 2000 |  | 2010 |  | 2020 |  | Single land use dynamic index/% |  |  |
| --- | --- | --- | --- | --- | --- | --- | --- | --- | --- |
|  | area/km<sup>2</sup> | ratio/% | area/km<sup>2</sup> | ration/% | area/km<sup>2</sup> | ration/% | 2000-2010 | 2010-2020 | 2000-2020 |
| Cultivated land (CL) | 1261.15 | 32.14 | 1125.85 | 28.70 | 1003.29 | 25.57 | -1.07 | -1.09 | -1.02 |
| Forest land (FL) | 2071.01 | 52.78 | 2054.28 | 52.36 | 2046.22 | 52.15 | -0.08 | -0.04 | -0.06 |
| Orchard land (OL) | 12.76 | 0.33 | 15.73 | 0.40 | 19.25 | 0.49 | 2.33 | 2.24 | 2.55 |
| Grassland (GL) | 309.00 | 7.88 | 263.56 | 6.72 | 266.23 | 6.79 | -1.47 | 0.10 | -0.69 |
| Water body (WB) | 105.33 | 2.68 | 114.44 | 2.92 | 115.74 | 2.95 | 0.87 | 0.11 | 0.49 |
| Built-up land (BL) | 109.15 | 2.78 | 245.61 | 6.26 | 335.65 | 8.55 | 12.50 | 3.67 | 10.37 |
| Unused land (UL) | 55.09 | 1.40 | 104.02 | 2.65 | 137.13 | 3.49 | 8.88 | 3.18 | 7.44 |
| total | 3923.49 | - | 3923.51 | - | 3923.49 | - | 21.96 | 8.17 | 19.08 |

**Fig 2.**
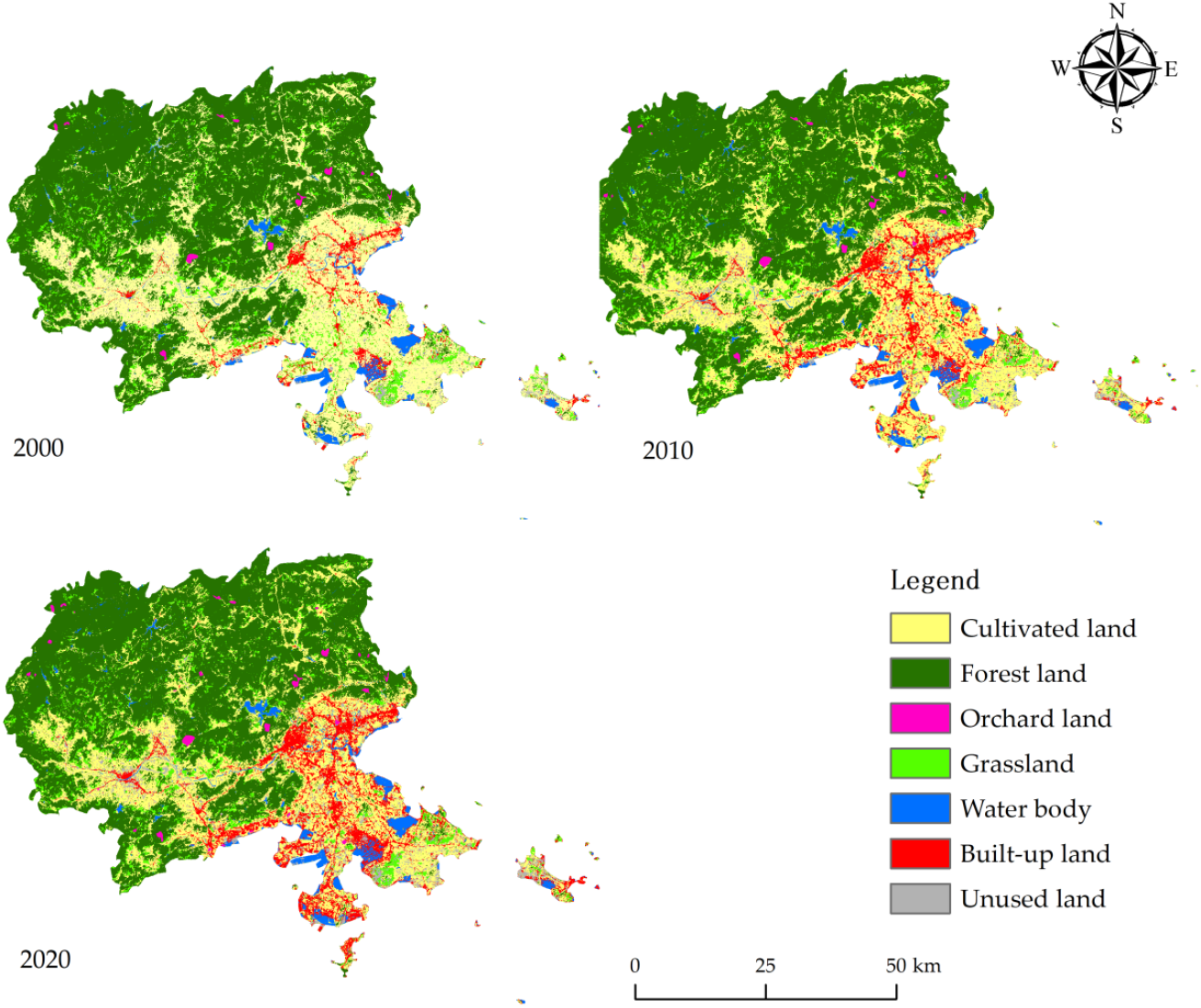
Current land-use in Putian city from 2000 to 2020.

The land use dynamic index can be also used to estimate the change of land use structure in a specific period from another aspect. The results in Table 1 showed that the average values of single land use dynamic index at the two stages of 2000-2010 and 2010-2020 were 21.96% and 8.17% respectively, implying that the land use transformation of Putian city had been continuous in the past 20 years. The rate of land use change from 2000 to 2020 reached 19.08%, which meant that the land use transformation of Putian city was strongly interfered by human activities. From 2000 to 2020, the dynamic degrees of cultivated land, forest land and grassland were -1.02%, -0.06% and -0.69%, respectively, indicating that the area of these land types showed a downward trend; the dynamic degree of the water body was 0.49%, performing a slow increasing trend; the dynamic degree of the orchard land was 2.55%, indicating that the increase rate was relatively fast; the dynamic degree of built-up land was the largest, reaching 10.37%; the dynamic degree of unused land was the second, with 7.44%, showing that the increase rates of built-up land and unused land were accelerating.

From the distribution results of land use (Fig 2), the study found that the types of land use change in the city were distinct in spatial distribution. From the center of the city to the edge were built-up land, cultivated land, forest land, etc. The cultivated land was mainly concentrated in Xiuyu district in the southeast of Putian city and Xianyou county in the southwest; forest land was mainly distributed in Hanjiang district in the northeast and Xianyou county in the northwest; the grassland was mainly distributed in the plain area in the southeast of Putian city, and the grassland area in Xiuyu district accounted for the largest proportion, which might be related to the conversion of deserted dry land to grassland; the orchard land was mainly distributed in the dry land and platform of Hanjiang district and Xianyou county, while the water body was mainly distributed in the coastal areas such as Hanjiang district and Xiuyu district, except for the reservoirs in the districts and counties. The fish farming in these areas was more developed than in inland areas.

### Tupu analysis of land use transformation

#### Tupu of land use transformation from 2000 to 2010

The spatial distribution of land-use transformation Tupu units in Putian city from 2000 to 2010 showed significant differences. The most obvious changes of Tupu units were “cultivated land→ built-up land”(code 16), “grassland → built-up land” (code 46) and “cultivated land → unused land”(code 17), accounting for 28.56%, 23.79% and 18.18% of the total converted land use types respectively (Table 2). The spatial distribution of these three types of Tupu units in the study area was a large-scale high-density distribution (Fig 3a). The Tupu units of “cultivated land→forest land “(code 12) and” forest land → cultivated land” (code 21) accounted for 6.70% and 6.03% of all converted land use types, respectively. The Tupu units of “cultivated land→water body” (code 15) and “cultivated land→grassland” (code 14) were also noteworthy, accounting for 4.11% and 3.34% of the total converted land use types, respectively. These land use types were widely distributed in the south and coastal areas of the study area.

**Table 2.**
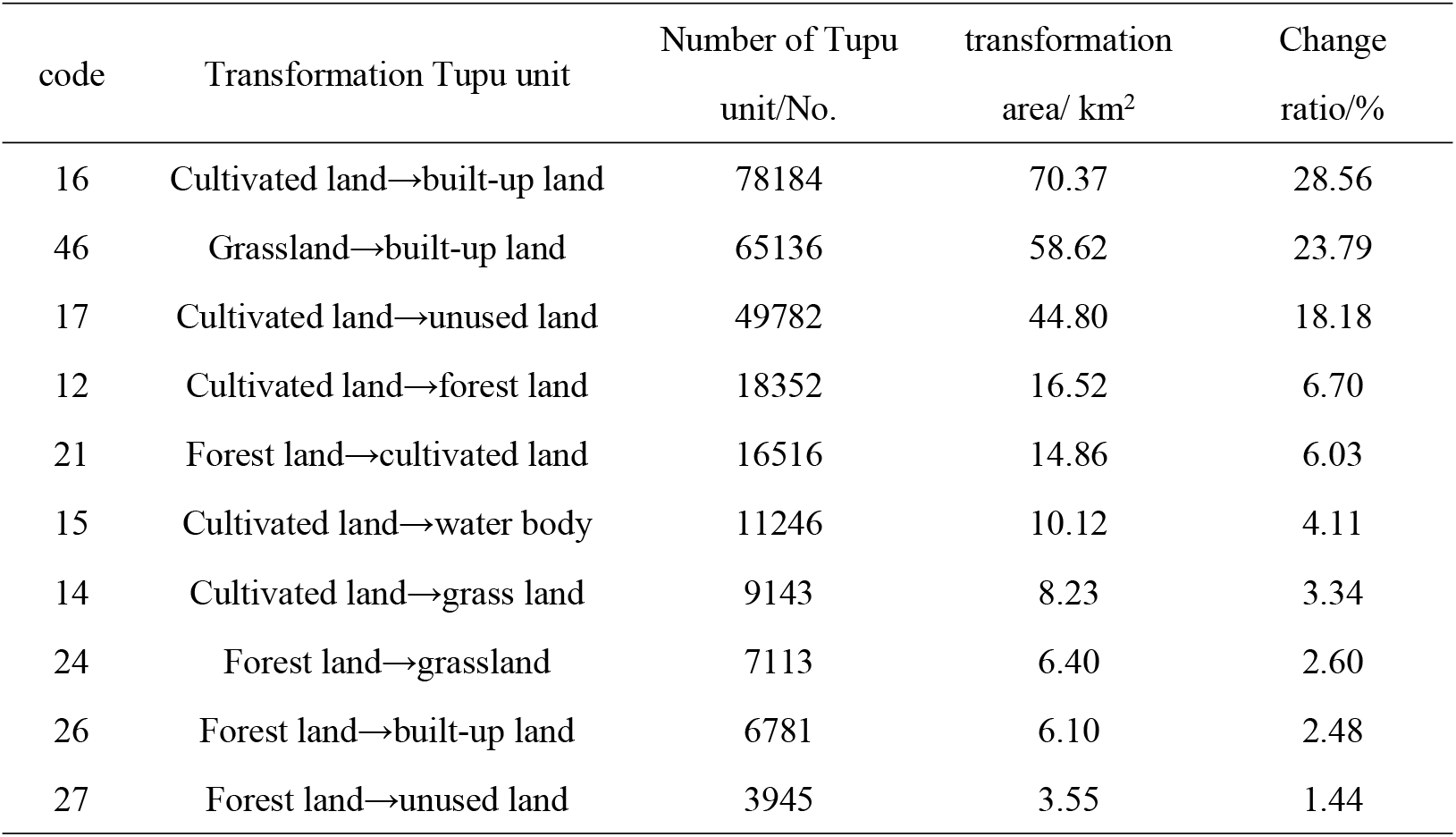
Tupu unit ranking of land-use transformation in Putian city from 2000 to 2010 (Top 10).

**Fig 3.**
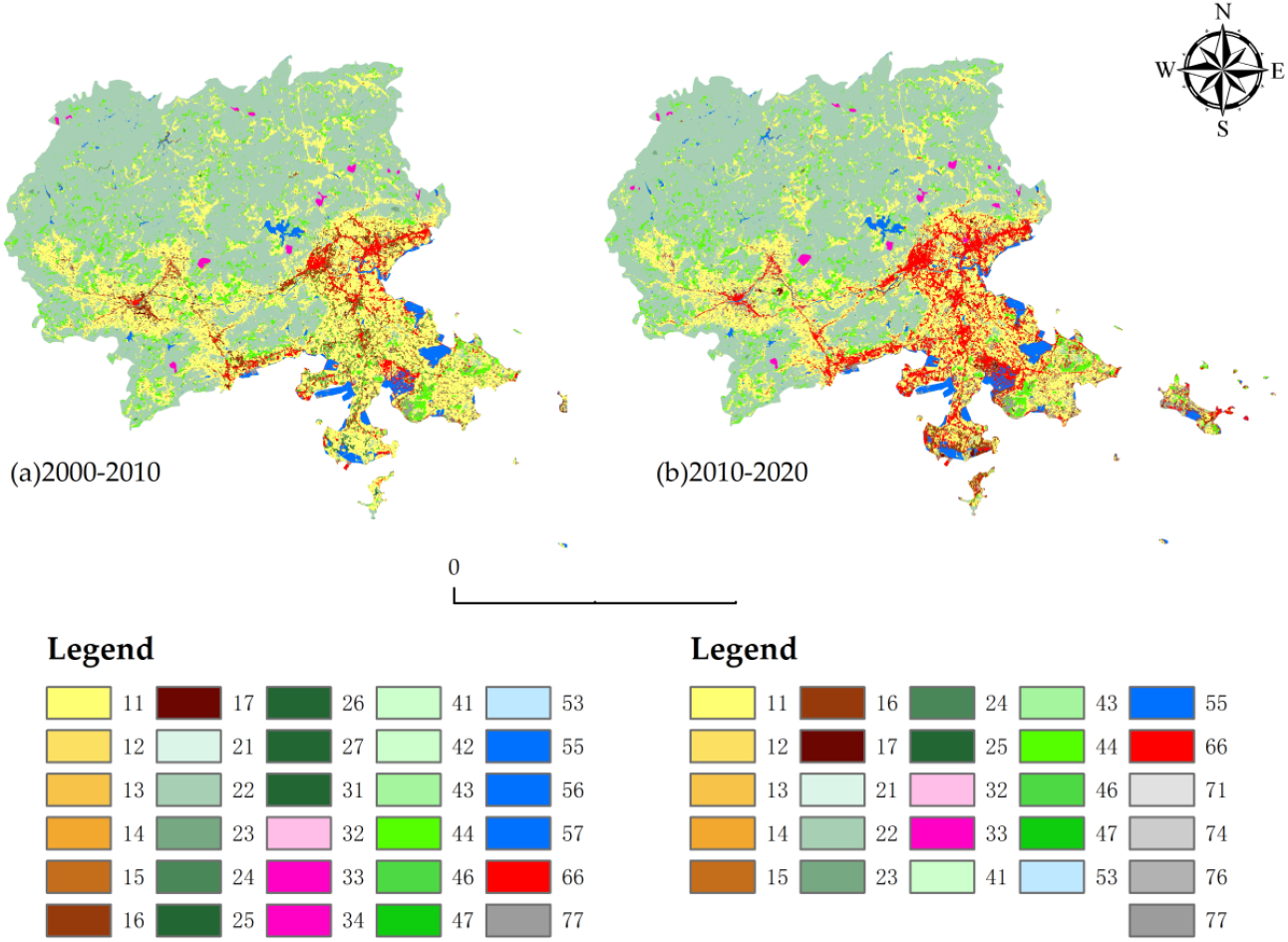
Tupu of land-use transformation in Putian city from 2000 to 2020.

Further from the transformation area of each land type (Table 2), the results showed that cultivated land was the type with the largest transfer-out area among all land types, up to 150.94 km^2^. 46.62% of the area was converted into built-up land (70.37 km^2^), 29.68% into unused land (44.80 km^2^), 10.94% into forest land (16.52 km^2^), 6.71% into water body (10.12 km^2^), 5.45% into grassland (8.23 km^2^) and 0.60% into orchard land (0.90 km^2^); The transfer-in area of cultivated land was only 15.63 km^2^, of which 95.10% was from forest land (14.86 km^2^) and the rest was from grassland (0.77 km^2^). From the perspective of the spatial distribution of cultivated land transferred out, it was mainly converted into built-up land in urban areas, and forest land and orchard land in some mountainous areas such as Xianyou county and Hanjiang district.

#### Tupu of land-use transformation from 2010 to 2020

The total area of land use transfer from 2010 to 2020 was 183.21 km^2^, which was 63.17 km^2^ less than the previous period (2000-2010). Among all the transformation Tupu, the most important land type transformation was still cultivated land (Table 3), with a total of 146.38 km^2^ transferred out, and more than half of the area was converted to built-up land, accounting for 58.41% (85.50 km^2^), mainly distributed in coastal areas, especially in Xiuyu district (Fig 3b). The transfer-in area of cultivated land was only 23.80 km^2^, mainly from forest land and grassland. The proportion of increase and decrease of cultivated land was obviously unbalanced. The transferred-out area of forest land was 22.26 km^2^, and the proportions converted to cultivated land, grassland, orchard land and water body were 74.10% (16.50 km^2^), 17.21% (3.83 km^2^), 8.14% (1.81 km^2^) and 0.55% (0.12 km^2^), respectively. The transferred area of forest land was 14.10 km^2^, mainly from cultivated land and orchard land. The transferred-out area of grassland (9.01 km^2^) was smaller than the transferred in area (7.73 km^2^), with a difference of 1.28 km^2^. The grass land transferred out mainly converted to cultivated land (5.35 km^2^), and the area transferred in mainly came from cultivated land (7.49 km^2^). The conversion of cultivated land to grassland was mainly due to abandoned cultivated land, which finally evolved into grassland, and most of the grassland was finally transferred into built-up land.

**Table 3.** Tupu unit ranking of land use transformation in Putian city from 2010 to 2020 (Top 10).

| code | Transformation Tupu unit | Number of Tupu<br>unit /No. | Transformation<br>area/ km<sup>2</sup> | Change ratio<br>/% |
| --- | --- | --- | --- | --- |
| 16 | Cultivated land→built-up land | 94999 | 85.50 | 46.67 |
| 17 | Cultivated land→unused land | 40640 | 36.58 | 19.96 |
| 21 | Forest land→cultivated land | 18329 | 16.50 | 9.00 |
| 12 | Cultivated land→forest land | 15545 | 13.99 | 7.64 |
| 14 | Cultivated land→grassland | 8320 | 7.49 | 4.09 |
| 41 | Grassland→cultivated land | 5939 | 5.35 | 2.92 |
| 24 | Forest land→grassland | 4256 | 3.83 | 2.09 |
| 76 | Unused land→built-up land | 3154 | 2.84 | 1.55 |
| 71 | Unused land→cultivated land | 2175 | 1.96 | 1.07 |
| 23 | Forest land→orchard land | 2014 | 1.81 | 0.99 |

### Analysis of land-use rising / falling Tupu

#### Land-use rising Tupu

The difference of land use structure transformation in Putian city was studied by generating the rising Tupu of Putian city in 2000-2010 and 2010-2020 (Fig 4) and making statistics and analysis on the transferred data (Table 4).

**Table 4.**
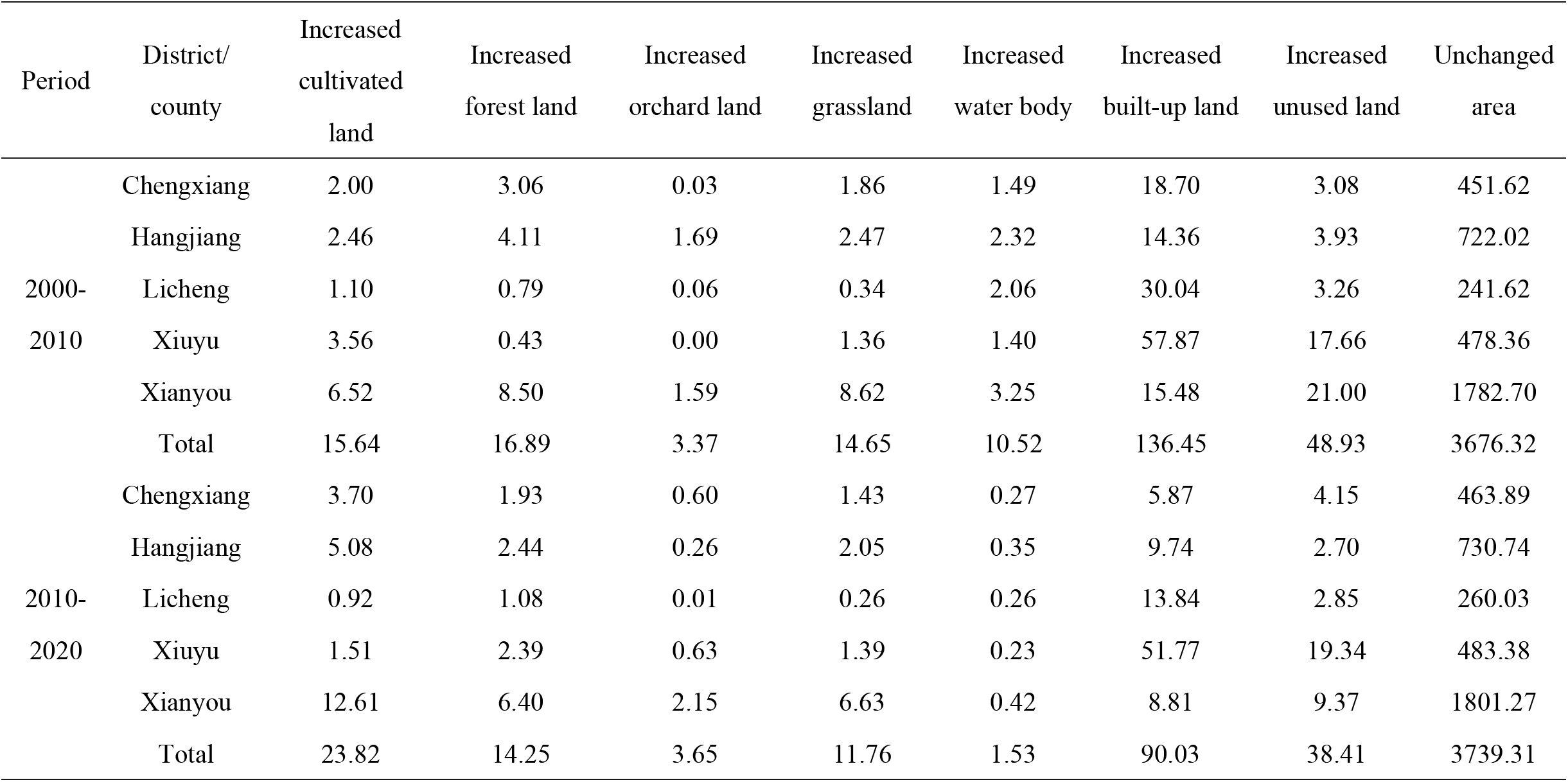
The structure list of land use rising Tupu in Putian city from 2000 to 2020 (km^2^).

| Period | District/<br>county | Increased<br>cultivated<br>land | Increased<br>forest land | Increased<br>orchard land | Increased<br>grassland | Increased<br>water body | Increased<br>built-up land | Increased<br>unused land | Unchanged<br>area |
| --- | --- | --- | --- | --- | --- | --- | --- | --- | --- |
| 2000-2010 | Chengxiang | 2.00 | 3.06 | 0.03 | 1.86 | 1.49 | 18.70 | 3.08 | 451.62 |
|  | Hangjiang | 2.46 | 4.11 | 1.69 | 2.47 | 2.32 | 14.36 | 3.93 | 722.02 |
|  | Licheng | 1.10 | 0.79 | 0.06 | 0.34 | 2.06 | 30.04 | 3.26 | 241.62 |
|  | Xiuyu | 3.56 | 0.43 | 0.00 | 1.36 | 1.40 | 57.87 | 17.66 | 478.36 |
|  | Xianyou | 6.52 | 8.50 | 1.59 | 8.62 | 3.25 | 15.48 | 21.00 | 1782.70 |
|  | Total | 15.64 | 16.89 | 3.37 | 14.65 | 10.52 | 136.45 | 48.93 | 3676.32 |
| 2010-2020 | Chengxiang | 3.70 | 1.93 | 0.60 | 1.43 | 0.27 | 5.87 | 4.15 | 463.89 |
|  | Hangjiang | 5.08 | 2.44 | 0.26 | 2.05 | 0.35 | 9.74 | 2.70 | 730.74 |
|  | Licheng | 0.92 | 1.08 | 0.01 | 0.26 | 0.26 | 13.84 | 2.85 | 260.03 |
|  | Xiuyu | 1.51 | 2.39 | 0.63 | 1.39 | 0.23 | 51.77 | 19.34 | 483.38 |
|  | Xianyou | 12.61 | 6.40 | 2.15 | 6.63 | 0.42 | 8.81 | 9.37 | 1801.27 |
|  | Total | 23.82 | 14.25 | 3.65 | 11.76 | 1.53 | 90.03 | 38.41 | 3739.31 |

**Fig 4.**
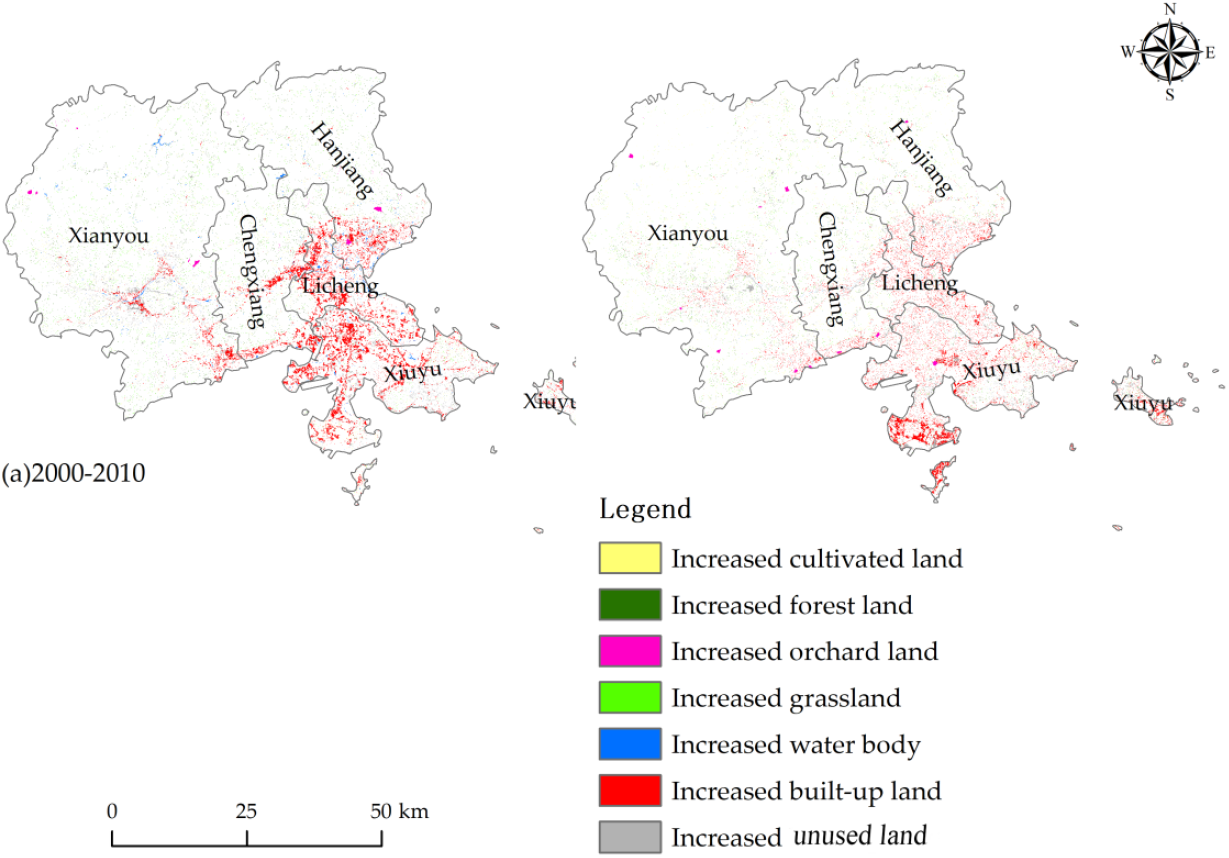
Land-use rising Tupu in Putian city from 2000 to 2020.

As shown in Table 4 and Fig 4 that from 2000 to 2010, the changed area and unchanged area of land use in Putian city were 246.45 km^2^ and 3676.32 km^2^, respectively, accounting for 6.28% and 93.72% of the area of the study area. Among the changed areas, the area of newly increased built-up land was the largest, reaching 136.45 km^2^, accounting for 55.37% of the total newly increased area, followed by the newly increased unused land (48.93 km^2^), the newly increased forest land (16.89 km^2^), the newly increased cultivated land (15.64 km^2^), the newly increased grassland (14.65 km^2^), the newly increased water body (10.52 km^2^) and the newly increased orchard land (3.37 km^2^). From the perspective of districts and counties, Xianyou county had the highest proportion of unchanged areas (48.49%), and the land use change was the most stable; The unchangeable area in Licheng district accounted for the lowest proportion (6.57%), and its newly added built-up land area was the highest at 30.04 km^2^. In terms of the area of newly added land types, Xianyou county had the largest area of newly added cultivated land at 6.52 km^2^, followed by Xiuyu district (3.56 km^2^), Hanjiang district (2.46 km^2^), Chengxiang district (2.00 km^2^), and Licheng district (1.10 km^2^). In terms of forest land, Xianyou county, Hanjiang district and Chengxiang district were the three areas with the largest newly added forest land, accounting for 92.78% of all newly added forest land. As the main fruit production bases in Putian city, Hanjiang district and Xianyou county had a newly added orchard land of 3.28 km^2^, accounting for 97.04% of the total newly added orchard land. It is noteworthy that among the newly added built-up land, Xiuyu district had the largest newly added area, reaching 57.87 km^2^, followed by Licheng district, Chengxiang district, Xianyou county and Hanjiang district. Therefore, the transfer of land use to built-up land was the most active in Xiuyu district during this period, while the Chengxiang district was relatively stable.

From 2010 to 2020, the area of land use change in Putian city decreased compared with the previous stage (2000-2010), with a total change area of 183.44 km^2^. The newly increased area of built-up land was 90.03 km^2^, with the highest rise. The area of newly increased cultivated land reached 23.82 km^2^, accounting for 12.99% of the total newly increased area; the area of newly increased forest land and grassland decreased by 2.64 km^2^ and 2.89 km^2^ respectively compared with the previous stage (2000-2010). From the perspective of districts and counties, the newly increased cultivated land area of Xianyou county was 12.61 km^2^, reaching the largest, followed by the newly increased area of 5.08 km^2^ in Hanjiang district. The area of newly increased built-up land in all districts and counties decreased significantly compared with the previous stage (2000-2010), among which the area of newly increased construction land in Licheng district had the largest reduction, with a reduction of 16.20 km^2^. In addition, except Licheng district and Xiuyu district, the area of newly increased forest land in other districts and counties had decreased compared with the previous stage, and Xianyou county had the largest reduction. The newly added orchard land was mainly allocated in Xianyou county (2.15 km^2^), Xiuyu district (0.63 km^2^), Chengxiang district (0.60 km^2^) and Licheng district (0.01 km^2^).

In general, the characteristics of land-use rise in Putian city showed high similarity in 2000-2010 and 2010-2020. It should be noted that the main newly increased land types of all districts and counties in the two periods were used for new construction land areas, but the increasing rate appeared to slow down in the later period.

#### Land use falling Tupu

The temporal and spatial evolution of land use change in Putian city was studied by generating the falling Tupu of Putian city in 2000-2010 and 2010-2020 (Fig 5) and making statistics and analysis on the transferred data (Table 5).

According to Table 5 and Fig 5, it indicated that the shrinking area of cultivated land in Putian city from 2000 to 2010 reached 150.94 km^2^, accounting for 61.24%, followed by the shrinking area of grassland and forest land, accounting for 24.38% and 13.64% respectively; the shrinking area of orchard land, water area and unused land was relatively small, accounting for less than 1%. From the perspective of occurrence area, the shrinkage of cultivated land was mainly concentrated in Xianyou county and Xiuyu district, accounting for 55.55%. The main types of shrinking land in the central area of Putian (Chengxiang district and Licheng district) were cultivated land and grassland (62.76 km^2^), much larger than the newly added area of the two land types (5.30 km^2^) in the same period. Therefore, the land use change in the entire urban areas was highlighted by the significant shrinking of cultivated land area and the sharp expansion of built-up land during this period.

From 2010 to 2020, the shrinking area and structure of land types in Putian city maintained a high similarity with the previous period. It was worth noting that although the shrinking area of cultivated land was not much different from the previous period, it was still the land type with the largest shrinking area, indicating that the trend of cultivated land occupation at this phase had not been reversed. Compared with the previous period (2000-2010), the grassland shrinking area in Xiuyu district decreased significantly, from 32.66 km^2^ in the previous period to 2.31 km^2^, with a decrease of 92.93%. The shrinking area of other land types was the smallest, and the change was not obvious. Among all districts and counties, Xiuyu district had the highest shrinking area of cultivated land, reaching 71.56 km^2^; followed by Xianyou county with the shrinking area of cultivated land at 27.65 km^2^. Other districts and counties had also decreased. The shrinking area of forest land and grassland in Xianyou county was 12.12 km^2^ and 4.46 km^2^ respectively. The shrinking area of forest land in Hanjiang district was 4.76 km^2^. In general, compared with the shrinking area of land use types in the previous stage, the reduced areas in this stage tended to be lower, but the spatial distribution had expanded.

#### Key driving force of land use change

Canonical correlation analysis showed that the total eigenvalue of the ranking axis was 0.9504 in terms of the correlation between the development of social and economic indicators (environmental factors) and the ranking axis, and the first two ranking axes explained 82.22% and 12.82% of the total data respectively. From Table 6 and Fig 6, it showed that the changes of 17 social and economic statistical indicators of Putian city and its districts and counties in the past 20 years had a great correlation with the first and second ranking axes. The fishery output value (l), the gross value of the primary industry (g), the gross domestic product (f), the total industrial output value (m), the urbanization rate (c), the gross value of the tertiary industry (I) and the gross value of the secondary industry (h) were significantly positively correlated with the first ranking axis, while the non-agricultural population (a), the total retail sales of social consumer goods (e), the total population (b), the total agricultural output value (J), and the forestry output value (k) were significantly negatively correlated with the first ranking axis. The second axis was negatively correlated with the economic crop area (q), agricultural population (d), pesticide use (o), grain crop area (P) and agricultural fertilizer application (n). Therefore, the first ranking axis could be regarded as the growth axis of socio-economic production input and output while the second one was mainly the growth axis of indicators related to agricultural economy, which would not be analyzed and explained because of low interpretation rate. It showed that more than 95% of the relationship between land use change and socio-economic development was reflected on the first ranking axis, which reflectes the major information about the relationship between socio-economic indicators and land-use change. Therefore, the results of canonical correspondence analysis could well explain the correlation between land use change and development in socio-economic statistical indicators.

**Table 6.** Correlation between socioeconomic indicators and ranking axis in Putian.

| Code | Indicators | CCA1 | CCA2 |
| --- | --- | --- | --- |
| l | Fishery output value | 0.373071 | 0.3432 |
| g | The gross value of the primary industry | 0.132691 | 0.432421 |
| f | The gross domestic product | 0.061692 | 0.662864 |
| m | The total industrial output value | 0.059091 | 0.700707 |
| c | The urbanization rate | 0.04841 | 0.214636 |
| i | The gross value of the tertiary industry | 0.043508 | 0.627488 |
| h | The gross value of the secondary industry | 0.036183 | 0.727325 |
| q | Economic crop area | -0.003031 | -0.335187 |
| a | The non-agricultural population | -0.020272 | 0.24007 |
| e | The total retail sales of social consumer goods | -0.04027 | 0.657806 |
| b | The total population | -0.074218 | 0.009524 |
| d | Agricultural population | -0.094883 | -0.062492 |
| o | Pesticide use | -0.121958 | -0.245523 |
| p | Grain crop area | -0.168553 | -0.457895 |
| j | The total agricultural output value | -0.196017 | 0.326368 |
| n | Agricultural fertilizer application | -0.268036 | -0.093573 |
| k | Forestry output value | -0.324892 | 0.649751 |

**Fig 6.**
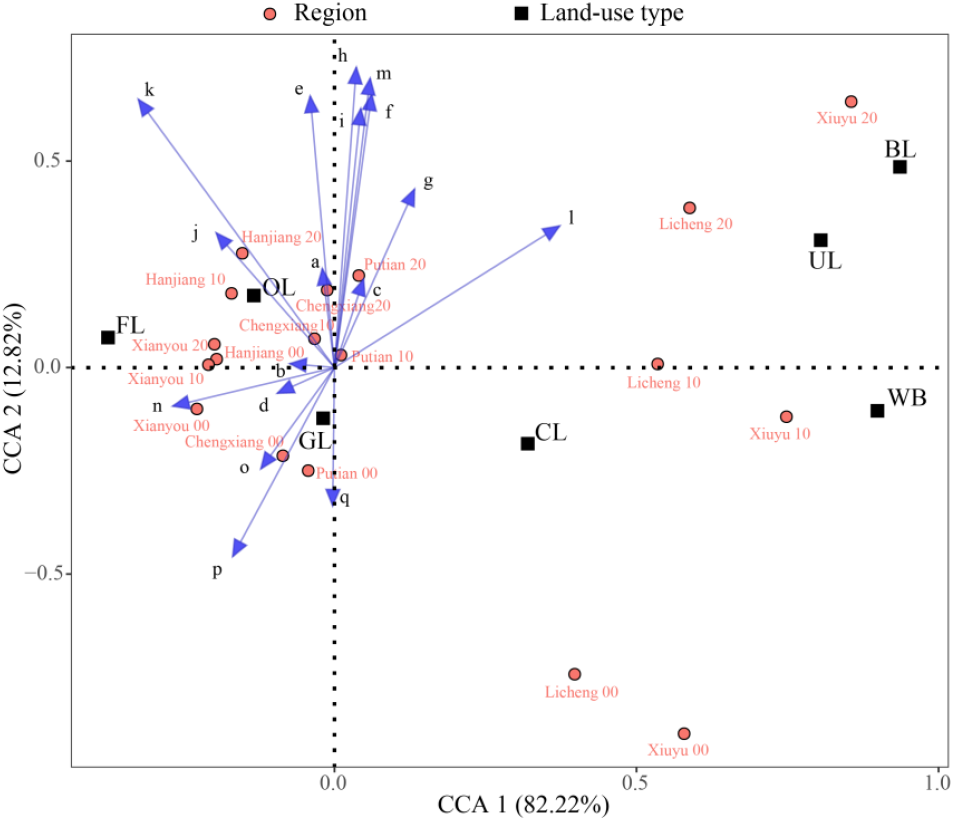
CCA 2-d ranking of the land use change and socioeconomic indicator change in districts or counties of Putian.

Abbreviations and letters are the same as those in Table1 and 6, respectively.

According to the importance ranking of the selected 17 socioeconomic indicators (Table 7), there were 6 indicators with extremely significant differences. The larger *r*^2^ and smaller *Pr* indicated the index was more important. The order of importance was the gross value of the secondary industry, the output value of forestry, the total industrial output value, the gross domestic product, the total retail sales of social consumer goods, and the gross value of the tertiary industry. In CCA ranking (Fig 6), built-up land, water body and unused land were located on the upper right side of the first principal axis, the cultivated land on the lower right side, the orchard land and forest land on the upper left side of the first principal axis, and the grassland on the lower left side. It indicated that the growth of social and economic production input and output drove the area increase of built-up land, resulting in the extrusion of cultivated land, grassland and forest land. This was closely related to the growth of the gross value of the secondary industry and the total industrial output value of Putian city by 13.09 and 13.16 times respectively in the 20 years from 2000 to 2020. At the same time, the built-up land increased by 2.07 times, while the cultivated land, grassland and forest land decreased by 20.45%, 13.84% and 1.97% respectively. According to the located quadrant of districts and counties, Xiuyu district and Licheng district were on the right side of the first main axis, positively correlated with built-up land and unused land (in fact, most of them were also planned to be built) and negatively correlated with cultivated land. It showed that in the past 20 years, the secondary industry in these two districts had developed rapidly, and the use of non-agricultural land had increased by 1.91 and 2.75 times respectively, resulting in a significant reduction in the cultivated land, decreasing by 24.79% and 31.00% respectively; at the same time, abandonment of agriculture for business increased the unused land by 2.67 and 1.10 times respectively. The quadrant location truly reflected the relative position and relationship between the changes of land use types and the development of major socioeconomic indicators in the districts and counties of Putian city, which clearly displays the driving effect of socioeconomic indicators on the change of land use type, which led to the corresponding change of ecosystem service values.

**Table 7.** Importance of socioeconomic indicators of Putian City.

| Code | Indicator | $r^2$ | $Pr (>r)$ | significant difference |
| --- | --- | --- | --- | --- |
| h | The gross value of the secondary industry | 0.5457 | 0.001 | *** |
| k | Forestry output value | 0.5295 | 0.008 | ** |
| m | The total industrial output value | 0.5121 | 0.002 | ** |
| f | The gross domestic product | 0.4603 | 0.003 | ** |
| e | The total retail sales of social consumer goods | 0.4455 | 0.008 | ** |
| i | The gross value of the tertiary industry | 0.4105 | 0.009 | ** |
| l | Fishery output value | 0.3092 | 0.054 |  |
| p | Grain crop area | 0.2496 | 0.109 |  |
| g | The gross value of the primary industry | 0.2359 | 0.125 |  |
| j | The total agricultural output value | 0.1342 | 0.341 |  |
| q | Economic crop area | 0.1265 | 0.352 | . |
| o | Pesticide use | 0.0704 | 0.567 |  |
| a | The non-agricultural population | 0.0590 | 0.643 |  |
| n | Agricultural fertilizer application | 0.0554 | 0.652 |  |
| c | The urbanization rate | 0.0547 | 0.679 |  |
| d | Agricultural population | 0.0035 | 0.976 |  |
| b | The total population | 0.0000 | 1.000 |  |

\*\* and \*\*\* refer to significant difference at 1% and 0.1% level.

## Discussion

Based on the land use data obtained from Landsat TM / OLI remote sensing images in 2000, 2010 and 2020, the study systematically analyzed the land use change characteristics of Putian city by using the methods of single land use dynamics and Geo-information Tupu, then further revealed the driving force of land use change in Putian city and its five districts and counties based on the results of CCA analysis. The results could be provided as references for optimizing regional land use pattern and improving regional ecological environment. The main research conclusions were as follows:

The land use pattern of Putian city was mainly composed of cultivated land and forest land and the rest of the land types were embedded in them. The built-up land had a trend of expanding outward. The study found that from 2000 to 2020 the areas of the cultivated land, forest land and grassland decreased year by year, while the orchard land, water body, built-up land and unused land increased year by year. Cultivated land was the land type with the largest decline among all land types and showed a continuous downward trend. Specifically, it decreased from 1261.15 km^2^ in 2000 to 1003.29 km^2^ in 2020, with a decrease of 20.45%; the grassland had the second largest reduction, with a decrease of 42.77 km^2^ in 20 years, and the forest land was the land with the smallest reduction in area among all land types. The area of built-up land increased year by year during the period of 2000-2020, which was reflected in the fact that the built-up land in Putian city in 2000 was only 109.15 km^2^, accounting for 2.78% of the total area, and had doubled to 335.65 km^2^ after 20 years of construction since Putian city was currently in the rapid development stage of urbanization and industrialization, in which the construction land increasingly demanded. Much ecological land, including cultivated land, had been requisitioned in the case, affecting the ecological environment quality of Putian city, and causing widespread concern in the society [39]. This resulted in the distinct spatial distribution in Putian city, indicating that the cultivated land was mainly concentrated in Xiuyu district in the southeast of Putian city and Xianyou county in the southwest. Historically, the main cultivated lands in Putian city were mainly distributed in the Xinghua Plain under the jurisdiction of Hanjiang and Licheng districts, the eastern and western plains of Xianyou county, the southern plains and the Fengjiang Plain. The plains were mostly below 60 meters above sea level. However, the urbanization development occupies too much plain farmland, which may leave behind farmland with relatively poor transportation and water conservancy, such as the hilly terraces in the southwest northwest mountains of Xianyou and the coastal dry land in the southeast Xiuyu district, which seriously affects the healthy development of agriculture. The forest land was mainly distributed in Hanjiang district in the northeast and Xianyou county in the northwest, which is not affected by urbanization because of inconvenient transportation and far away from cities and towns; In addition, in recent years in order to protect the mother river Mulan River in Putian city, the government has strengthened the construction and protection of the ecological forest at the source of Mulan River in the northeast of Hanjiang district and the northwest of Xianyou county[40]. The grassland was mainly distributed in the plain area in the southeast of the city, and the grassland area of Xiuyu district accounted for the largest proportion. Since Xiuyu district as a new district, has accelerated its urbanization, much construction land to be approved for construction is overgrown with weeds, resulting in an increase in grassland; moreover, the district is located in the coastal dry land, with serious disrepair of water conservancy and low crop yield and benefit, forcing local farmers to abandon agriculture for business, resulting in the outflow of labor force, which leads to the increase of abandoned land, and then turns into grassland with weeds spreading The orchard land was mainly distributed in Hanjiang district and Xianyou county. This is because Putian city is located in the middle subtropical region, which is suitable for the development of regional characteristic fruit trees. Hanjiang district and Xianyou county attach importance to the development of subtropical fruit trees, so they are the main producing areas of litchi, longan, loquat, pomelo, etc. The water body was mainly distributed in the coastal areas of Hanjiang, Xiuyu and Licheng districts, in addition to the reservoirs in the districts and counties.

The spatial distribution of Tupu units of land use transformation in Putian city showed significant differences, and the area of cultivated land and grassland transferred-out was the most obvious. The study found that from 2000 to 2020, other land types in Putian city were transferred into built-up land via the different routes. A total of 226.48 km^2^ was transferred into built-up land in 20 years, mainly in the economically developed areas around the cities and counties, especially Xiuyu district. The main types of transferred Tupu unit in 2000-2010 and 2010-2020 were “cultivated land → built-up land” and “cultivated land → grassland → built-up land”. In the past 20 years, 297.32 km^2^ of cultivated land had been transferred out and 39.43 km^2^ had been transferred in. The proportion of increase and decrease of cultivated land was obviously unbalanced. Among them, the expansion rate of built-up land in Xiuyu district was the fastest among all districts and counties, with an increase rate of 275%, and a total expansion of 109.75 km^2^ in 20 years. Under the premise of limited land resources, the continuous expansion of urban land certainly would occupy other land types, especially production and ecological lands. In addition to the contraction of cultivated land caused by the expansion of built-up land, various areas of Putian city had been abandoned to varying degrees and finally evolved into deserted field(grassland), resulting in waste of limited land resources due to poor agricultural production conditions (such as saline alkali land), low grain efficiency, rural labor transfer and other factors. Putian city was the main grain sales area in Fujian Province, and its grain supply was highly dependent on external sources. Its basic farmland was mainly distributed in Xiuyu district and Xianyou county. Thus, Putian city faced great challenges in ensuring food security and curbing the “non-agricultural” and “non grain” of cultivated land. Therefore, the government should attach great importance to such phenomenon. We believed that in addition to optimize the layout of urban, rural, industrial and mining land and carry out the renovation of abandoned land, it is important to improve the utilization rate of land resources. For example, the research team found that the fields in Xiuyu district of Putian city were mostly saline and alkaline land with poor fertility, and most of them could only grow sweet potatoes, soybeans and other crops. What is more, the monoculture practice led to continuous cropping obstacles, resulting in a decline in yield and quality [39,41]. Taking Dongzhuang Town, Xiuyu district, Putian city as a pilot, the research team proposed the strategy of effectively utilizing the space-time resources of cultivated land and combining with agriculture, culture and tourism, that is, planting famous, special and excellent crops with less investment, short planting period and high efficiency, implementing rotation and intercropping system to improve the multiple cropping index and expanding agricultural functions by holding farmers harvest festival, research and learning experience and other projects. In the practice, we had helped the local farmers build a large-scale modern agricultural demonstration field in the local area, create an ecological leisure agricultural brand, and turn the saline-alkali land that was not suitable for the growth of crops into “golden land”, which had increased the income of farmers, consolidated and expanded the achievements of poverty alleviation, won the recognition of the local government, new business entities and farmers, and further promoted its application.

The CCA analysis showed that the macro-economic development objectives in a specific period, especially the urban expansion and the development of the secondary industry caused by the merger of cities and counties, were the key factors driving the spatial and temporal evolution of land use and the differential distribution pattern in Putian city. After 2002, Putian city and county merged to form Chengxiang district, Licheng district, Hanjiang district, Xiuyu district, Meizhou Island Management Committee and Meizhou Bay North Bank Management Committee, which inevitably promoted urban expansion, and objectively led to the increase of non-agricultural land and the reduction of cultivated land. Data showed that from 2000 to 2020, the area of newly increased built-up land in Putian city was the largest, reaching 226.49 km^2^, accounting for 52.68% of the total newly increased area while the shrinking area of cultivated land with 297.32 km^2^ was the largest, accounting for 69.15% of the total shrinking area. This could also be verified from the overall layout of “two points, three lines and three levels” in Putian City Master Plan (1993-2010). According to the plan, Putian strived to realize the common development of the north and south triangle areas, and the overall spatial layout of Chengxiang district and Hanjiang district had gradually formed a pattern of “cluster structure and zoning balance” [42]. During the same period, the development of Meizhou Bay Area in the south of Putian (including the districts of Xiuyu and Licheng, and two administrative committees) had become an important direction for the development of Putian city. In addition, according to the investigation, the Fujian Provincial Party Committee and the Provincial Government issued the notice on the overall plan for the comprehensive reform experiment of urban-rural integration in Putian city (MWF [2012] No. 5) in July 2012, which officially approved Putian city as a pilot project for the comprehensive reform of province’s urban-rural integration. In 2014, Putian city was approved as the first batch of new urbanization comprehensive reform pilot in China. Putian city took the urbanization of people as the core, and gradually explored and formed the Putian local urbanization mode of “five integration and five transformation”. Therefore, the urbanization rate increased after 2015 and the Putian on-site urbanization mode of “five integration and five modernization” had been gradually explored and formed. In 2020, the urbanization rate of the whole city rose to 62.7%, promoting the national economy and the per capita economic disposable amount. Those historical events as mentioned above further certified our results and explained in part the reason of land use change in the past 20 years.

Therefore, it is foreseeable that the city’s non-agricultural land would inevitably continue to grow, causing a series of environmental and ecological economic problems. At the same time, the transfer of population, the increase of non-agricultural population and the low agricultural efficiency had also accelerated the process of reducing the area of cultivated land. In addition, leaving farming for trade had caused serious farmland abandoned. It therefore is extremely urgent to carry out purposeful utilization and planning of land space, especially agricultural ecological transformation in the efficient use of cultivated land.

It was also worth noting that the newly added forest land and orchard land mainly occurred in Hanjiang district and Xianyou county, which could be attributed to Putian city’s in-depth implementation of afforestation projects, the policy of returning farmland to forests and orchard land, and the drive of comparative interests. Part of the abandoned farmland and slope cropland had been transformed into economic forests and ecological forests, creating a new pattern of forest and fruit industry with the common development of bionic cultivation of Chinese medicinal plants under the forest, loquat, Wendan pomelo, etc. We believe that under the current policy constraints of strictly controlling the occupation of ecological protection red line and permanent basic farmland, it is an inevitable choice to efficiently utilize land resources and develop forest economy (such as medicinal plants, forest tourism and other modes) in mountainous areas that are not easily accessible to the industrial chain, which can strive for greater space for regional sustainable development and ecological balance. At present, the authors strongly suggested that attention should be paid to the construction of the ecological source, especially in seven mountainous villages and towns of northwest Xianyou County and ecological barrier and ecological corridor construction for the coastal upland of southeast of Putian city. At the same time, with the aid of rural revitalization in China and Mulan river comprehensive treatment engineering in Putian, it is needed to do a good job of restoration and protection of water conservancy and farmland soil and effectively promote the ecological transformation of agricultural land to improve agricultural comprehensive effect.

## Conclusions

In short, in the past 20 years, due to the division of districts and counties in Putian city, urban expansion and urbanization have been significantly accelerated. This is driving significant changes in land use pattern, leading to the decrease of cultivated land and the increase of construction land. The results truthfully reflected the overall situation of land use change in Putian city during the period of 2000 -2020, indicating that the area of forest land occupied the largest proportion, followed by that of cultivated land, cultivated land and grassland, which decreased by 20.45% 1.20% and 13.83%, respectively. Among all land categories, cultivated land is the land category with the largest area decline and shows a continuous decreasing trend, while woodland is a relatively stable land category, and the area of construction land and unused land is increasing year by year. In terms of the land distribution characteristics of different regions, the cultivated land after the change was mainly concentrated in the coastal dry upland of Xiuyu district in the southeast and the plain farmland of Xianyou county in the southwest of Putian city. The forestland is mainly distributed in the northeast of Hanjiang district and the northwest of Xianyou county. Grassland is mainly distributed in the plain area in the southeast of the city, and Xiuyu district has the largest proportion of grassland area. The garden area is mainly distributed in Hanjiang district and Xianyou county and other places; Waters mainly distributed in Hanjiang River and Xiuyu district coastal area. From the perspective of the area transferred by different types of land, arable land is the largest type of land transferred, among which 46.62% is converted to construction land, 29.68% to unused land, 10.94% to woodland, 6.71% to water body, 5.45% to grassland, and 0.60% to garden land. In terms of the spatial distribution of cultivated land transfer, it is mainly converted to construction land in urban areas, and converted to woodland and garden land in some mountainous areas such as Xianyou county and Hanjiang district. The results truly reflect the geographical, ecological and economic characteristics of Putian, the coastal city, which will provide a research basis for further exploring the optimization strategy for land use of Putian city.

## Acknowledgments

We would like to thank all the authors who contributed papers to this collection. We would also like to thank the PLOS ONE staff for their valuable support.

## Reference

1. Long HL, Qu Y. Land use transitions and land management: A mutual feed back perspective. Land Use Policy. 2018; 74: 111–120. doi: 10.1016/j.landusepol.2017.03.021.

2. Wei J, Liu LL, Wang HY, Zhang YX, Wang CL, Liu JT, et al. Spatiotemporal patterns of land-use change in the Taihang Mountain (1990-2020). Chinese Journal of Eco-Agriculture. 2022; 30(7): 1123–1133. doi: 10.12357/cjea.20210870

3. Chen WX, Chi GQ, Li JF. The spatial association of ecosystem services with land use and land cover change at the county level in China, 1995-2015. Science of The Total Environment. 2019; 669(1): 459–470. doi: 10.1016/j.scitotenv.2019.03.139.

4. Su K, Wei DZ, Lin WX. Evaluation of ecosystem services value and its implications for policy making in China-A case study of Fujian Province. Ecological Indicators. 2020; 108: 105752. doi: 10.1016/j.ecolind.2019.105752.

5. Wu D, Li H, Ai N, Huang T, Gu JS. Predicting spatiotemporal changes in land use and habitat quality based on CA-Markov: A case study in central Ningxia, China. Chinese Journal of Eco-Agriculture. 2020; 28(12): 1969–19 78. doi: 10.13930/j.cnki.cjea.200221

6. MA (Millennium Ecosystem Assessment). Ecosystems and human well-being: a framework for assessment. Washington DC: Island Press; 2005.

7. Houghton RA. Terrestrial fluxes of carbon in GCP carbon budgets. Global Change Biology. 2020; 26(5): 3006–3014. doi: 10.1111/gcb.15050.

8. Arneth A, Barbosa H, Benton T, et al. Climate change and land: summary for policymakers. An IPCC Special Report on Climate Change, Desertification, Land Degradation, Sustainable Land Management, Food Security, and Greenhouse Gas Fluxes in Terrestrial Ecosystems, 2019, 43.

9. Su RQ, Cao YG, Wang WX, Qiu M, Song L. Analysis of spatiotemporal c haracteristics of cultivated land use change from 2001 to 2017 in the Cha obai River basin of the Beijing-Tianjin-Hebei region. Journal of Agricultur al Resources and Environment. 2020; 37(4): 574–582. doi: 10.13254/j.jare.2019.0266.

10. Wang ZC, Gao ZQ. Analysis on spatiotemporal characteristics and causes of tidal flat wetland in Jiaozhou bay from 1987 to 2017 based on land us e change. Research of Soil and Water Conservation. 2020; 27(6): 196–201. doi: 10.13869/j.cnki.rswc.2020.06.027.

11. Tong XR, Yang QY, Bi GH. Spatio-temporal characteristics of land use changes in Chongqing during 2000-2015. Resources and Environment in the Yangtze Basin. 2018; 27(11): 2481–2495. doi: 10.11870/cjlyzyyhj201811010.

12. Yu YH, Li ZJ, Lin JK, Liu JY, Wang S. TUPU characteristics of spatiotemporal variation for land use in the Yihe River Basin. Journal of Natural Resources. 2019; 34(5): 975–988. doi: 10.31497/zrzyxb.20190506.

13. Zhang Y, Peng JD, Wang JJ, Yang H. Analysis on spatial and temporal c hange of land use in Jianghan Plain based on Geo-information atlas. Rese arch of Soil and Water Conservation. 2020; 27(4): 85–92. doi: 10.13869/j.c nki.rswc.2020.04.012. doi: 10.13869/j.cnki.rswc.2020.04.012.

14. Cui XP, Guo YX. Analysis on the spatio-temporal dynamic evolution of la nd use structure of western urban agglomerations in the past 40 years. Jou rnal of Arid Land Resources and Environment. 2022; 36(2): 16–24. doi: 10.13448/j.cnki.jalre.2022.030.

15. Chen WX, Zeng J. Decoupling analysis of land use intensity and ecosyste m services intensity in China. Journal of Natural Resources. 2021; 36(11): 2853–2864. doi: 10.31497/zrzyxb.20211110.

16. Yang QQ, Xu GL, Li AJ, Liu YT, Hu CS. Evaluation and trade-off of ec osystem services in the Qingyijiang River Basin. Acta Ecologica Sinica. 2021; 41(23): 9315–9327. doi: 10.5846/stxb202009032293.

17. Gu RH, Zhu YL. Impact mechanism of regional land use on the ecological optimization: a case study of Jiangsu Province. Economic Geography. 2021; 41(11): 201–208. doi: 10.15957/j.cnki.jjdl.2021.11.022.

18. Huang MQ, Li YB, Ran CH, Li MZ. Dynamic changes and transitions of agricultural landscape patterns in mountainous areas: A case study from the hinterland of the Three Gorges Reservoir Area. Journal of Geographical Sciences. 2021; 76(11): 2749–2764. doi: 10.1007/s11442-022-1984-7.

19. Zhang J, Zhu WB, Wu SY, Li SC. Simulation of temporal and special land use changes in Jing-Jin-Ji urban agglomeration using CLUE-S model. Acta Scientiarum Naturalium Universitatis Pekinensis. 2018; 54(1): 115–124. doi: 10.13209/j.0479-8023.2017.137.

20. Chen SP, Yue TX. Studies on Geo-Informatic Tupu and its application. Geographical Research. 2000; 19(4): 337–343.

21. Wang S, Zhang LL, Lin WB, Huang QS, Song YX, Ye M. Study on veg etation coverage and land-use change of Guangdong Province based on M ODIS-NDVI. Acta Ecologica Sinica. 2022; 42(6): 2149–2163. doi: 10.5846/stxb202104261100.

22. Du GM, Zhang R, Yu FR. Analysis of cropping pattern in black soil region of Northeast China based on geo-information Tupu. Chinese Journal of Applied Ecology. 2022; 33(3): 694–702. doi: 10.13287/j.1001-9332.202203.017.

23. Zhang WH, Lyu X, Shi YY, Sun PL, Zhang YS. Graphic characteristics of land use transition in the Yellow River Basin. China Land Science. 2020; 34(8): 80–88. doi: 10.11994/zgtdkx.20200731.204521.

24. Lu C, Zhou H, Zhang F, Dong GL, Fu JS. Land spatial transformation analysis in Shandong Province based on geo Information map. Transactions of the Chinese Society for Agricultural Machinery. 2021; 52(7): 222–230. doi: 10.6041/j.issn.1000-1298.2021.07.023.

25. Yu X, Pu LJ, Xu Y, Zhu M. Analysis of land use changes in relation to environmental variables in coastal city in Jiangsu Province from 1980 to 2010: a case study in Dongtai City. Resources and Environment in the Yangtze Basin. 2016; 25(4): 537–543. doi: 10.11870/cjlyzyyhj201604001.

26. Su K, Wei DZ, Lin WX. Regional differences and spatial patterns of urbanization efficiency in Fujian Province, China. Acta Ecologica Sinica. 2019; 39(15): 5450–5459. doi: 10.5846/stxb201808261819.

27. Breiman L. Random forests. Machine learning. 2001; 45: 5-32. doi. org/10.1023/A:1010933404324.

28. Jin YH, Liu XP, Chen YM, Liang X. Land-cover mapping using random forest classification and incorporating NDVI time-series and texture: a case study of central Shandong. International Journal of Remote Sensing. 2018; 39(23): 8703–8723. doi:10.1080/01431161.2018.1490976.

29. Wang LJ, Kong YR, Yang XD, Xu Y, Liang L, Wang SG. Classification of land use in farming areas based on feature optimization random forest algorithm. Transactions of the Chinese Society of Agricultural Engineering. 2020;36: 244–250. doi: 10.11975/j.issn.1002-6819.2020.04.029.

30. Becker WR, Ló TB, Johann JA, Mercante E. Statistical features for land use and land cover classification in Google Earth Engine. Remote Sensing Applications: Society and Environment. 2021; 21, 100459. doi: 10.1016/j.rsase.2020.100459.

31. Liu J, Liu JK, An JJ, Zhang C. Precise crop classification based on multi-features from time-series Landsat 8 OLI images and Random Forest Algorithm. Agricultural Research in the Arid Areas. 2020; 38(3): 281-288, 298. doi: 10.7606/j.issn.1000-7601.2020.03.37.

32. Matthiasa B, Martinb H. Mapping imperviousness using NDVI and linear spectral unmixing of ASTER data in the Cologne-Bonn region (Germany. Proc Spie, 2004, 12: 274–284.

33. Zha Y, Gao J, Ni S. Use of normalized difference built-up index in automatically mapping urban areas from TM imagery. International Journal of Remote Sensing. 2003; 24(3): 583–594. doi: 10.1080/01431160304987.

34. Liu HQ, Huete A. A feedback-based modification of the NDVI to minimize canopy background and atmospheric noise. IEEE Transactions on Geoscience and Remote Sensing. 1995; 33(2): 457–465. doi: 10.1109/TGRS.1995.8746027.

35. Tang CC, Li YP. Geo-information Tupu process of land use/cover change in polycentric urban agglomeration: A case study of Changsha-Zhuzhou-Xiangtan urban agglomeration[J]. Geographical Research. 2020; 39(11): 2626–2641. doi: 10.11821/dlyj020200207.

36. Wu LL, Li XY, Mao DH, Wang ZM. Urban land use classification based on remote sensing and multi-source geographic data. Remote Sensing for Natural Resources. 2022; 34(1): 127–134. doi: 10.6046/zrzyyg.2021061.

37. Kong XL, Wang KL, Chen HS, Zhang JG, Zhang MY. Canonical correspondence analysis of land-use change and socio-economic development in Hechi Prefecture, Guangxi Province. Journal of Natural Resources, 2007; 22(1): 131–140. doi: 10.11849/zrzyxb.2007.01.016.

38. Ter braak Cjf. Canonical correspondence analysis: a new eigenvector technique for multivariate direct gradient analysis. Ecology. 1986; 67:1169–1179. doi: 10.2307/1938672.

39. Wu HM, Wu LK, Wang JY, Zhu Q, Xu JH, Zheng CL, et al. The mecha nisms of the rhizosphere management on the remission in consecutive mon oculture problem and the improvement of soil quality. Ecological Science. 2016; 35(5): 225–232. doi: 10.14108/j.cnki.1008-8873.2016.05.031.

40. Zheng JY. On Xi Jinping’s idea and style of governing the country——from the perspective of the governance of Mulan River in Putian, Fujian. Journal of Hunan University of Science and Engineering. 2019; 8: 66–69. doi: 10.16336/j.cnki.cn43-1459/z.2019.08.017.

41. Weng PY, Zheng HY. Causes and mechanism on crop Continuous monoculture problems and its control strategy. Subtropical Plant Science. 2020; 49(2): 157–162. doi: 10.3969/j.issn.1009-7791.2020.02.014.

42. Wang YX, Wang YF, Zhang JW, Wang Q. Land use transition in coastal area and its associated eco-environmental effect: A case study of coastal area in Fujian Province. Acta Scientiae Circumstantiae. 2021; 41(10): 3927–3937. doi: 10.13671/j.hjkxxb.2021.0294.

